# Transverse Aortic COnstriction Multi-omics Analysis (TACOMA) uncovers pathophysiological cardiac molecular mechanisms

**DOI:** 10.1101/2024.01.26.577333

**Authors:** Enio Gjerga, Matthias Dewenter, Thiago Britto-Borges, Johannes Grosso, Frank Stein, Jessica Eschenbach, Mandy Rettel, Johannes Backs, Christoph Dieterich

## Abstract

Time-course multi-omics data of a murine model of progressive heart failure induced by transverse aortic constriction (TAC) provide insights into the molecular mechanisms that are causatively involved in contractile failure and structural cardiac remodelling. We employ Illumina-based transcriptomics, Nanopore sequencing, and mass spectrometry-based proteomics on samples from the left ventricle (LV) and right ventricle (RV, RNA only) of the heart at 1, 7, 21, and 56 days following TAC and Sham surgery. Here, we present TACOMA, as an interactive web-application that integrates and visualizes transcriptomics and proteomics data collected in a TAC time-course experiment. TACOMA enables users to visualize the expression profile of known and novel genes and protein products thereof. Importantly, we capture alternative splicing events by assessing differential transcript and exon usage as well. Co-expression-based clustering algorithms and functional enrichment analysis revealed overrepresented annotations of biological processes and molecular functions at the protein and gene levels. To enhance data integration, TACOMA synchronizes transcriptomics and proteomics profiles, enabling cross-omics comparisons. With TACOMA (https://shiny.dieterichlab.org/app/tacoma), we offer a rich web-based resource to uncover molecular events and biological processes implicated in contractile failure and cardiac hypertrophy. For example, we highlight: (i) changes in metabolic genes and proteins in the time course of hypertrophic growth and contractile impairment; (ii) identification of RNA splicing changes in the expression of Tpm2 isoforms between RV and LV; and (iii) novel transcripts and genes likely contributing to the pathogenesis of heart failure. We plan to extend these data with additional environmental and genetic models of heart failure to decipher common and distinct molecular changes in heart diseases of different aetiologies.

**Database URL:** https://shiny.dieterichlab.org/app/tacoma

## Introduction

### Background

Transverse Aortic Constriction (TAC), is a commonly used experimental technique to study the pathophysiological mechanisms of heart failure (HF). TAC involves the partial occlusion of the transverse aorta (mainly in mice) leading to pressure overload-induced cardiac hypertrophy and HF in the end. The TAC-induced adverse effects typically depend on the degree of the aorta constriction as well as its duration (1). Over time, the TAC-induced pressure overload causes progressive remodelling of the heart in both left and right ventricles (LV and RV) and some of such responses include changes in gene expression, inflammatory responses, fibrosis, etc. (2). In this context, TAC has been established in animal models to understand the dynamic changes in molecular mechanisms associated with the transition from compensatory hypertrophy to heart failure. Multi-omics integration in the context of transverse aortic constriction can provide a comprehensive understanding of disease mechanisms, leading to potential insights into therapeutic strategies. Time-course multi-omics studies are particularly suitable for TAC since they allow the identification of dynamic changes and temporal patterns of key molecules involved in disease progression (1). Such an approach would then allow us to identify early molecular markers that precede HF as well as to propose potential therapeutic approaches.

In this study, we present a comprehensive multi-omics analysis based on a murine TAC model to study molecular changes during the progression of pressure overload-induced cardiac hypertrophy to heart failure (Figure 1). The study involved the collection of samples from the left ventricle (LV) and right ventricle (RV) of the heart at 1, 7, 21, and 56 days following TAC and Sham surgery. Besides TAC and Sham, we additionally have measurements from healthy mouse tissues at time-point 0 days, referred to as the Control samples. To comprehensively analyse the multi-layered molecular landscape of hypertrophy progression, we employed three distinct omics techniques: Illumina RNA-seq and Nanopore sequencing (long read cDNA sequencing) as well as proteomics (LV only). All were produced in triplicates. The Nanopore cDNA data was used to reconstruct a *de novo* assembled transcriptome. This allows us to identify novel transcript isoforms and provide a more comprehensive view of the alternative splicing landscape and isoform switching dynamics during TAC progression. Furthermore, proteomics analysis was only performed on the LV samples due to input material limitations. Proteomics is complementary to RNA-seq approaches as it targets another molecular layer of TAC-induced hypertrophy across different time points.

**Figure 1:**
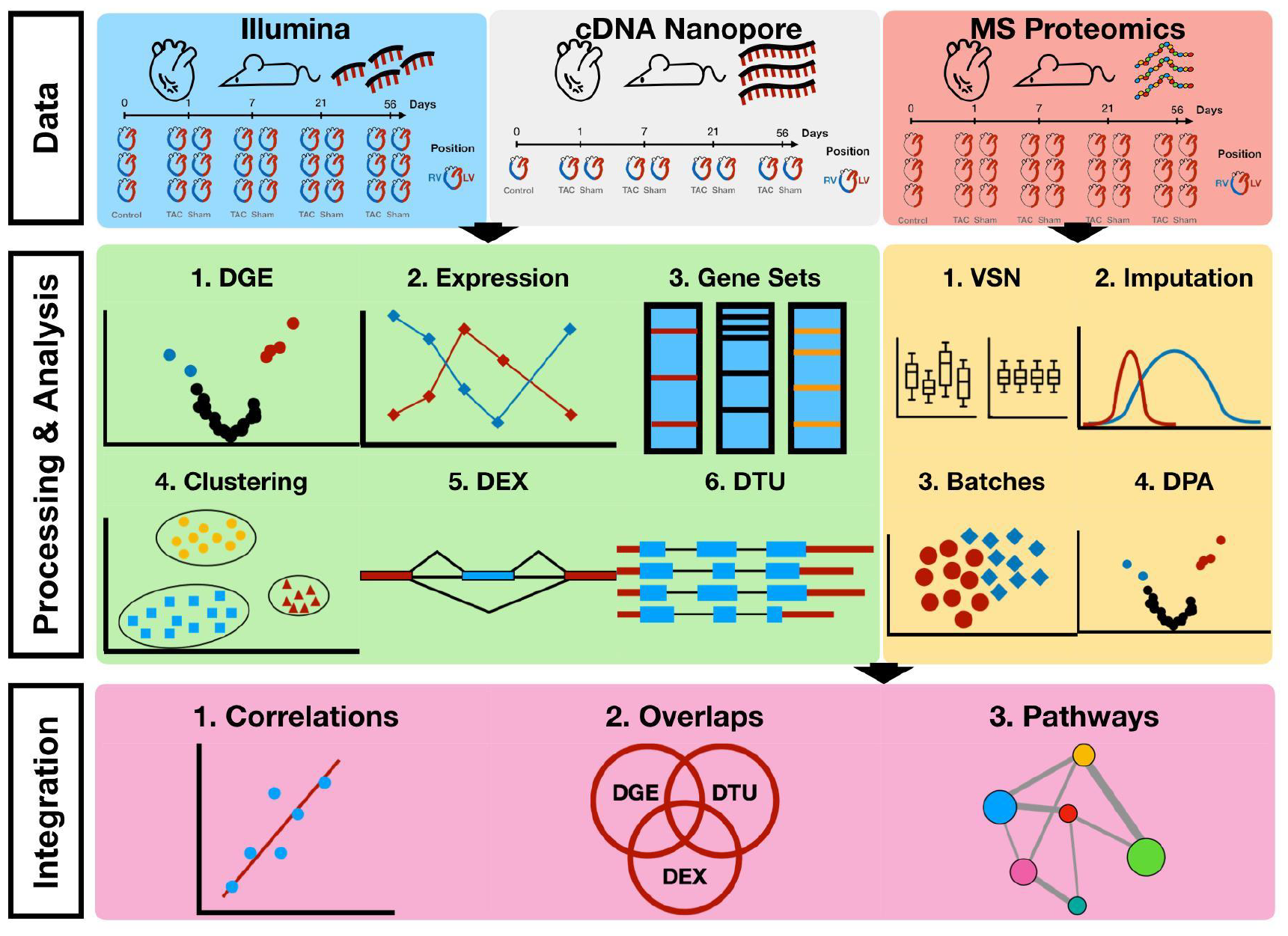
Workflow of multi-omics data analysis in TAC mouse model. Data consists of Illumina and Nanopore cDNA transcriptomics of TAC, Sham and Control (RV and LV) as well as Mass-spectrometry Proteomics (LV only) at time-points 0, Day 1, Day7, Day 21 and Day 56. Differential gene/transcript/exon expression/usage analyses were performed over the transcriptomics data along with enrichment and clustering analyses. Differential protein abundances were estimated from raw mass-spectrometry proteomics upon processing of the data (normalisation, imputation, batch effect correction) and enrichment analyses were performed. Cross-omics integration of transcriptomics and proteomics allowed for a comparison between the two data modalities.

To advance the visualisation, interpretation and accessibility of our new data, we have developed TACOMA as an online application, which allows any user to investigate the molecular mechanisms behind HF progression. TACOMA provides functionalities which allow the visualisation of analysis results with a special focus on differential gene expression (DGE), function enrichment analysis, gene co-expression modules, differential exon and transcript usage analysis (DXE and DTU) as well as differential protein abundances. To the best of our knowledge, TACOMA is the first interactive web application to encompass such a wide range of analyses in the context of TAC and cardiomyopathy progression (Figure 2).

**Figure 2:**
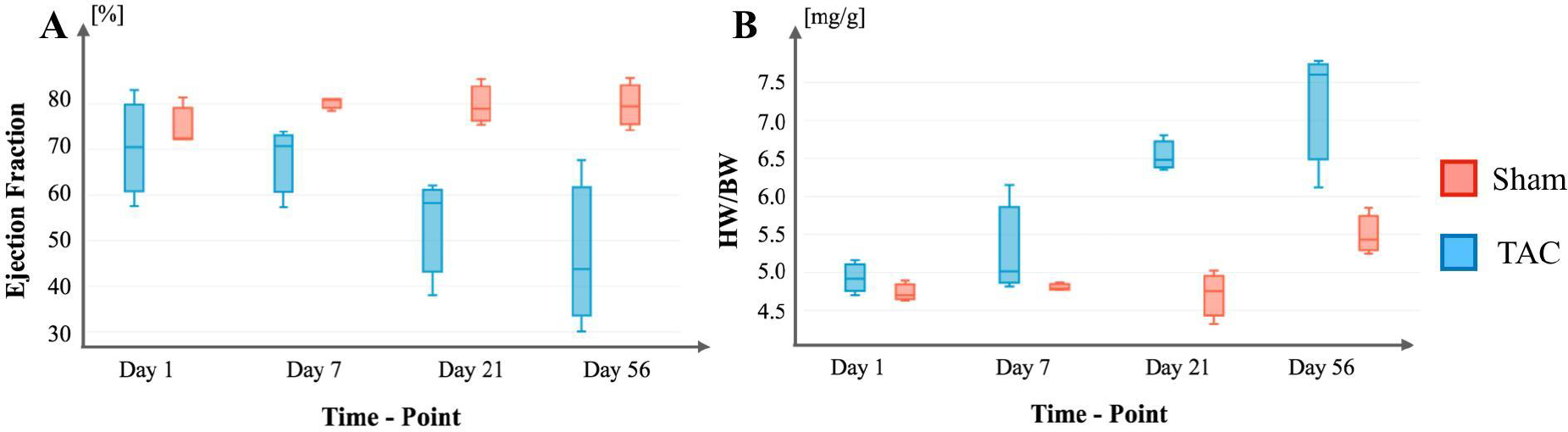
HF progression in the described TAC mouse model. **A)** Progression of Ejection Fraction (EF) changes across time in TAC and Sham conditions. **B)** Progression of Heart-Weight to Body-Weight (HW/BW) ratios across time in TAC and Sham conditions.

## Methods

### Sequencing data and processing

#### Illumina total RNA-seq

Total RNA from left and right ventricular tissue were rRNA depleted and subject to stranded RNA-seq library preparation for the Illumina platform at the Cologne Center for Genomics (CCG). All libraries were sequenced on a NovaSeq 6000 in paired-end mode (2x 100bp) at an average depth of 50 million fragments per library.

#### Nanopore cDNA sequencing

Total RNA from left and right ventricular tissues were polyA-selected and subject to cDNA library preparation on the ONT Nanopore platform (Kit: SQK-DCS109 and Flow cell: FLO-MIN106). All libraries were sequenced on an ONT GridION X5 device at an average depth of 1 million long reads per library.

#### Illumina read processing, mapping and counting

We first removed adaptors and low-quality bases with Flexbar (v3.5.0) (3). We then identified reads that aligned to mouse tRNA or rRNA sequences using Bowtie2 (v2.3.5.1) (4) and discarded them. The remaining reads were aligned to the mouse EnsEMBL 102 genome with STAR (2.6.0c) (5). We observed an average proportion of unique mapping reads above 70% throughout all libraries. We generated gene, transcript and exon count tables using StringTie2 (v2.1.3b) (6) and ballgown (2.28.0) (http://bioconductor.org/packages/ballgown/).

#### Nanopore processing and mapping

Nanopore base calling was performed with Guppy 3.4.5 using the dna_r9.4.1_450bps_hac.cfg model. Nanopore reads were mapped against the EnsEMBL 102 reference genome with minimap 2.22 (7).

#### Guided transcriptome assembly

The transcription assembly was performed using StringTie2 (v.2.1.7) on autosomes and sex chromosomes, and features from other chromosomal regions were discarded. First, we paired a cDNA library that had been sequenced with ONT and Illumina and executed StringTie2 in the guided mode, using Ensembl as a reference. Next, we applied StringTie2 to each individual library, using the merged annotation obtained from the first step as a guide. Finally, the annotations were merged to create a unified annotation. We merged transcript annotation with the StringTie2 merge command and removed transcript isoforms representing less than 10% of relative transcript abundance or having less than 3 reads. The reference gene and transcript names as well as the class codes were obtained by running GffCompare (v0.12.2) (8) against the reference annotation GRCm38.102. Upon the transcriptome assembly, transcript counts were quantified with salmon (v1.10.1) (9) and gene counts were obtained from transcript counts by using the *DESeqDataSetFromTximport()* function from DESeq2 R-package (v.140.2) (10).

### Analysis of Gene Expression Data

Differential Gene Expression (DGE) analysis was performed with edgeR (v3.38.4) (11). The analysis accounted for multiple variables, including Condition (TAC & Sham), Ventricle (RV & LV) as well as Time/Day (Day 0, 1, 7, 21 & 56). A full model design matrix was formulated, consisting of the ventricle, day, condition, interaction between ventricle and condition, and the interaction between day and condition as follows: model.matrix(∼ Ventricle + Day + Condition + Ventricle:Condition + Day:Condition, data=metatada). In the design matrix, the interaction terms capture the combined effects of Condition and Ventricle, as well as Condition and Day, on gene expression. The interaction term between Condition and Ventricle accounts for how the gene expression is influenced by TAC/Sham depending on the sample position (RV or LV). Similarly, the interaction term between Condition and Day reveals whether the effect of TAC/Sham on gene expression varies with the specific day of measurement. By including these interaction terms in the design matrix, we aim to better capture the relationships between the variables and potentially identify dependent effects among them. The details of the ‘metadata’ table used in the design matrix can be found in the Supplementary Materials (Supplementary Table 1). To guide the selection of appropriate comparisons for our DGE analysis, several key questions were considered:

i. which genes change by condition (TAC/Sham) globally?
ii. which genes change by ventricle (LV/RV) globally?
iii. which genes change in TAC for a day?
iv. which genes change at all time-points compared to control?
v. which genes change over time?
vi. which genes change on any day?
vii. which genes change in TAC for ventricle?

Considering the above, we have performed DGE analyses for a total of 12 comparisons and details about each (design and description) have been provided in the Supplementary (Supplementary Notes 1).

### Differential Exon and Transcript Usage Analysis

Differential exon and transcript usage analyses (DEX or DTU) were conducted to examine the variability of alternative splicing across different conditions or comparisons. First, we compute exon and transcript counts for every expressed gene using ballgown. Then, for example, testing for differential exon usage is equivalent to testing whether the exons in each gene have the same log-fold-changes as the other exons in the same gene. To perform DEX analysis, we utilised the edgeR v3.38.4 R-package (11), which allowed us to identify exons with differential expression in the same comparisons as those used for differential gene expression (DGE) analysis. Gene annotations were fetched from the Ensembl BioMart database (version November 2020) using the biomaRt v2.54.0 R-package (12), to associate gene symbols and descriptions with Ensembl gene IDs. Exon counts were filtered to only include entries with a maximum unique read count greater than 20 across all samples. The exon counts were then normalised by using the *calcNormFactors()* function and a generalised linear model was fitted to the data using the *glmFit()* function from edgeR. The differential exon usage was tested for the 12 comparisons by using the *diffSpliceDGE()* function and the results were then filtered to identify genes with significant differential exon usage at an FDR threshold of 0.05 according to the Simes method. The same strategy was used to test for DTU.

### Analysis of Novel Genes

The assembled sequences were scanned for open reading frames (ORFs) using ORFik (v1.20.2) (13) to identify potential coding sequences within the novel gene transcripts. The predicted ORFs were then translated into protein sequences via Biostrings (v2.68.1). The translated sequences were then used for the subsequent domain annotation step with Interproscan (14) to provide insights into the potential functions of the novel proteins. The domain annotations included the domain IDs from various primary databases (PFAM, PantherDB, CATH-Gene3D), along with the signature name and description of each domain, offering detailed insights into the protein characteristics.

### Gene Co-expression Networks

To identify groups of genes with similar co-expression patterns, we have followed a similar strategy as described in (15). To summarise, we have employed the WGCNA v1.72.1 R package (16) to analyse gene co-expression from RNA-seq data, focusing on 13,197 genes that met the criteria of being significantly regulated (adjusted p-value <= 0.05) in at least one of the 12 DGE comparisons. By using topological overlap (TO), genes were clustered to spot co-expression patterns. To ensure the reproducibility and robustness of clusters, a bootstrap resampling was performed and final co-expression modules were identified using hierarchical clustering, and their significance was validated through posthoc resampling and a Z-test. Associations between co-expressed gene networks and observed phenotypes were determined by calculating the biweight midcorrelation between genes and biological traits or disease association for continuous physiological variables (i.e. Ejection Fraction - Supplementary Table 2). For binary/discrete variable correlation such as the pathological (TACvsSham), positional (RVvsLV) and temporal levels (Day7vsDay1, Day21vsDay7, Day56vsDay21, etc.), the standard Pearson correlation was used instead of the biweight midcorrelation.

### Sample preparation for Proteomics

Reduction of disulphide bridges in cysteine-containing proteins was performed with dithiothreitol (56°C, 30 min, 10 mM in 50 mM HEPES, pH 8.5). Reduced cysteines were alkylated with 2-chloroacetamide (room temperature, in the dark, 30 min, 20 mM in 50 mM HEPES, pH 8.5). Samples were prepared using the SP3 protocol (17, 18) and trypsin (sequencing grade, Promega) was added in an enzyme-to-protein ratio 1:50 for overnight digestion at 37°C. Next day, peptide recovery in HEPES buffer by collecting supernatant on magnet and combining with second elution wash of beads with HEPES buffer. Peptides were labelled with TMT10plex (19) Isobaric Label Reagent (ThermoFisher) according to the manufacturer’s instructions. Samples were combined for the TMT10plex and for further sample clean up an OASIS® HLB μElution Plate (Waters) was used. Offline high pH reverse phase fractionation was carried out on an Agilent 1200 Infinity high-performance liquid chromatography system, equipped with a Gemini C18 column (3 μm, 110 Å, 100 x 1.0 mm, Phenomenex) (20).

### LC-MS/MS data acquisition

An UltiMate 3000 RSLC nano LC system (Dionex) fitted with a trapping cartridge (μ-Precolumn C18 PepMap 100, 5μm, 300 μm i.d. x 5 mm, 100 Å) and an analytical column (nanoEase™ M/Z HSS T3 column 75 μm x 250 mm C18, 1.8 μm, 100 Å, Waters). Trapping was carried out with a constant flow of trapping solution (0.05% trifluoroacetic acid in water) at 30 μL/min onto the trapping column for 6 minutes. Subsequently, peptides were eluted via the analytical column running solvent A (0.1% formic acid in water, 3% DMSO) with a constant flow of 0.3 μL/min, with increasing percentage of solvent B (0.1% formic acid in acetonitrile, 3% DMSO). The outlet of the analytical column was coupled directly to an Orbitrap Fusion™ Lumos™ Tribrid™ Mass Spectrometer (Thermo) using the Nanospray Flex™ ion source in positive ion mode. The peptides were introduced into the Fusion Lumos via a Pico-Tip Emitter 360 μm OD x 20 μm ID; 10 μm tip (New Objectives) and an applied spray voltage of 2.4 kV. The capillary temperature was set at 275°C. Full mass scan was acquired with mass range 375-1500 m/z in profile mode in the orbitrap with resolution of 120000. The filling time was set at a maximum of 50 ms with a limitation of 4x10^5^ ions. Data dependent acquisition (DDA) was performed with the resolution of the Orbitrap set to 30000, with a fill time of 94 ms and a limitation of 1x10^5^ ions. A normalized collision energy of 38 was applied. MS^2^ data was acquired in profile mode.

### Proteomics Database search

IsobarQuant (21) and Mascot (v2.2.07) were used to process the acquired data, which was searched against a customized database containing common contaminants and reversed sequences. The following modifications were included into the search parameters: Carbamidomethyl (C) and TMT10 (K) (fixed modification), Acetyl (Protein N-term), Oxidation (M) and TMT10 (N-term) (variable modifications). For the full scan (MS1) a mass error tolerance of 10 ppm and for MS/MS (MS2) spectra of 0.02 Da was set. Further parameters were set: Trypsin as protease with an allowance of maximum two missed cleavages: a minimum peptide length of seven amino acids; at least two unique peptides were required for a protein identification. The false discovery rate on peptide and protein level was set to 0.01.

### Analysis of Protein Abundance Data

### Data Preparation

Protein abundance data was prepared in a matrix format for analysis using the DEP2 v0.4.8.24 R-Package (22). The data was organised to include protein intensity levels quantified across different samples or conditions. For the processing and analysis of the protein abundance data, we have relied on an established workflow based on the DEP2 v0.4.8.24 R-Package (22).

#### Normalization

Protein intensities were normalised to mitigate the technical biases and variability. In this case, the normalisation step was performed by using the Variance Stabilization Normalization (VSN) method through the use of the *normalize_vsn()* function from the DEP2 package.

#### Imputation of missing values

As 16.85% of the data in our protein intensity matrix is missing, a data imputation strategy from the DEP2 package was employed to estimate the missing values. In this case, we have assumed that missing values originated from low-abundant proteins. Therefore, a strategy was employed to impute the missing data by filling it with random values generated from a Gaussian distribution centred around the lower 1% value of the distribution of existing data through the use of the *impute()* function from DEP2.

#### Batch effect correction

Principal Component Analysis (PCA) of the normalised and imputed data set revealed clustering of samples based on each replicate, thus suggesting the presence of batch effects which needed to be corrected. Technical variations associated with the observed batch effects were identified and removed from the dataset by applying the *removeBatchEffect()* function from the limma R-package (23).

#### Differential Protein Analysis (DPA)

DPA analyses were subsequently performed following batch effect correction to identify proteins exhibiting significant changes in abundance for the TAC vs Control, Sham vs Control and TAC vs Sham comparisons for all the time-points combined as well as at each time-point separately. For this, we have used the *test_diff()* function from DEP2 as it performs a differential enrichment test based on protein-wise linear models and empirical Bayes statistics using limma. False discovery rates were estimated by using fdrtool (24) with three adjustment methods: Benjamini-Hochberg, Strimmer’s and Storey’s q-values.

### GO Enrichment Analysis

We have conducted Gene Ontology (GO) term enrichment analysis for each of the DGE and DPA comparisons for the Biological Process (BP), Molecular Function (MF) and Cellular Components (CC) ontologies. Integrated functional term enrichment analysis (of genes with adjusted p-values <= 0.05), as well as visualization, was performed by using the CellPlot R-package (https://github.com/dieterich-lab/CellPlot).

### Over-Representation Analysis

Over-representation analysis (ORA) over gene sets has been performed by using the *fora()* function from the fgsea v1.22.0 R-package (25). ORA was performed to identify which Pathway and Hallmark sets (26) were enriched for each cluster obtained from the gene co-expression network analysis.

### TACOMA

We introduce TACOMA (https://shiny.dieterichlab.org/app/tacoma), an interactive web-based tool designed to explore molecular signatures of TAC. TACOMA visualises the above-mentioned analyses. The deployment strategy involves ShinyProxy and an internal PostgreSQL database. We conceived TACOMA as an easily navigable dashboard, intentionally designed to cater to a diverse audience of biomedical scientists delving into the molecular underpinnings of heart disease progression. Similar to Magnetique (27), we integrated an interactive tour outlining the functions of each module and the available options within the application. TACOMA provides detailed and interactive results for ten views:

#### Phenotype View

Provides a table with phenotype information about each mouse sample undergone transcriptomics analysis such as Condition (TAC or Sham), Ventricle (RV or LV), Day (0, 1, 7, 21 and 56), Ejection Fraction (EF) as well as Heart Weight to Body Weight ratios (HW.BW).

#### Expression Profile

Provides a time course view on gene and protein expression. A cross-reference to their corresponding EnsEMBL (ensembl.org) web page is provided whenever applicable (for the GRCm38.102 reference genome). Users may select each gene to display the normalised expression profiles (mean expression and the standard deviation) across all time points on gene and protein levels as well as a heatmap of Z-scaled expressions at the protein (LV only) and gene level (LV and RV).

#### Gene View

Provides results from the DGE analysis: i) a table of genes sorted from the most to the least significant (based on adjusted p-values); and ii) a volcano plot visualising the direction, magnitude and significance of changes in gene expression. On the sidebar users can select the question that they are interested in as well as the exact comparison that they wish to visualise as described in the Methods section. Additionally, the users can select from the table a desired gene to visualise as box-plots its CPM expression in groups of samples tailored to the selected comparison.

#### Proteomics View

Provides results from the DPA analysis: i) a table of proteins sorted from the most to the least significant (based on adjusted p-values); and ii) a volcano plot visualising the direction, magnitude and significance of changes in protein abundances. On the sidebar users can select the main comparison that they are interested in (TAC vs Control/Day0, Sham vs Control/Day0 or TAC vs Sham for all combined samples or for each time-point separately). Additionally, the users can select from the table a desired gene to visualise its normalised abundance values in groups of samples tailored to the selected comparison. Significantly regulated proteins have been highlighted in red in the volcano plots and the users can select from three adjustment methods that have been applied (Benjamini-Hochberg, Strimmer’s or Storey’s adjustment).

#### Gene Set View

Provides a tabulated representation of enriched Gene Ontology terms specific to each chosen DGE and DPA comparison sorted from the most to the least significantly enriched set. Gene-set enrichments were performed for three types of ontologies: Biological Process (BP), Molecular Function (MF) or Cellular Component (CC). After choosing a gene set, users can visualise the differential gene/protein expression/abundance of its members as well as their normalised expression across each sample grouped by their aetiology.

#### WGCNA View

Provides a tabulated list of groups of modules of co-expressed genes based on the WGCNA strategy described previously. This view additionally provides a heatmap which shows the correlation of each gene cluster to a specific phenotype as well as the significance of such correlation (with a cut-off p-value of 0.05). Once a gene cluster gets selected, users receive a dot plot for Pathway (Reactome and BIOCARTA) and Hallmark sets from MSigDB (26) enrichment (with adj-pvalue<=0.05).

#### DEX View

Provides a tabulated list of genes with differential exon usage sorted from the most to the least significant adjusted p-value score for a selected gene comparison. Gene structures are visualised through a ggtranscript plot (28), which shows the location and expressions of each exon. Significant exons (adjusted p-value <= 0.05) are highlighted in green in the top half of the exon depiction, while the level of its regulation (LogFC) is depicted in the bottom half (blue for up-regulation and red for down-regulation). This is complemented by a tabular representation below that contains the same exon-level information.

#### DTU View

Provides a tabulated list of differential transcript abundance values and their significance for a given gene comparison. Users may select a transcript from the table to visualise transcript proportions of the corresponding gene.

#### Integration View

Provides cross-omics comparison functionality between the proteomics and transcriptomics data for TAC vs Sham data at each time-point (LV only) as well as for all the time-points combined. This view provides a tabulated list of genes which appear to be significant in either DGE and DPA comparisons, or only in DGE’s or DPA’s (adjusted p-value <= 0.05) or in neither. Additionally, a scatter plot shows LogFC values of the gene or protein level of analysis. Finally, we provide gene set enrichment information on Pathway and Hallmarks sets (adjusted p-value <= 0.05) based on differential expression data from the two modalities.

#### Novel Genes

Provides an interface to display key information of novel genes i.e. not overlapping any known gene locus, which are significantly regulated (adjusted p-value <= 0.05) in at least one of the 12 DGE comparisons. A table provides a list of gene symbols that are novel; comparisons in which such a gene becomes significant; ID’s of its transcripts as well as the number of exons and transcripts that are part of such gene. Similar to DEX View, structures of novel genes can be visualized through ggtranscript upon the selection of a desired gene ID. Additionally, after the selection of a specific gene, additional detailed information will be displayed in a tabulated format, such as i) the DNA and protein sequences of the predicted ORFs; ii) the domain IDs associated with each sequence; iii) the signature name and iv) description for each predicted domain (when applicable). Lastly, the genomic context can be studied by using links to the respective locus in the EnsEMBL genome browser.

## Results

We demonstrate the utility of TACOMA by providing insights into potential biological processes that could be associated with the progression of cardiomyopathies.

### Enrichment of Oxidative Phosphorylation and Fatty Acid Metabolism Hallmarks

From the cross-omics comparison of DGE and DPA in the Integration View of TACOMA, we were able to identify the most significant enrichment at both proteomics and transcriptomics levels (LV only) for the combined TAC vs Sham comparison: ‘fatty acid metabolism’ and ‘oxidative phosphorylation’ (Figure 3A). Alterations in myocardial metabolism are a hallmark of heart failure, with a multitude of studies showing decreased cardiac mitochondrial ATP production, reduced TCA cycle flux and decreased fatty acid beta-oxidation in preclinical models and humans (29, 30). Herein, we focus on the time-dependent enrichment of the above-mentioned gene sets over time in the left ventricle (Supplementary Figures 1-6 for visualization of genes Figure 3, A-C for visualization of enrichment scores).

**Figure 3:**
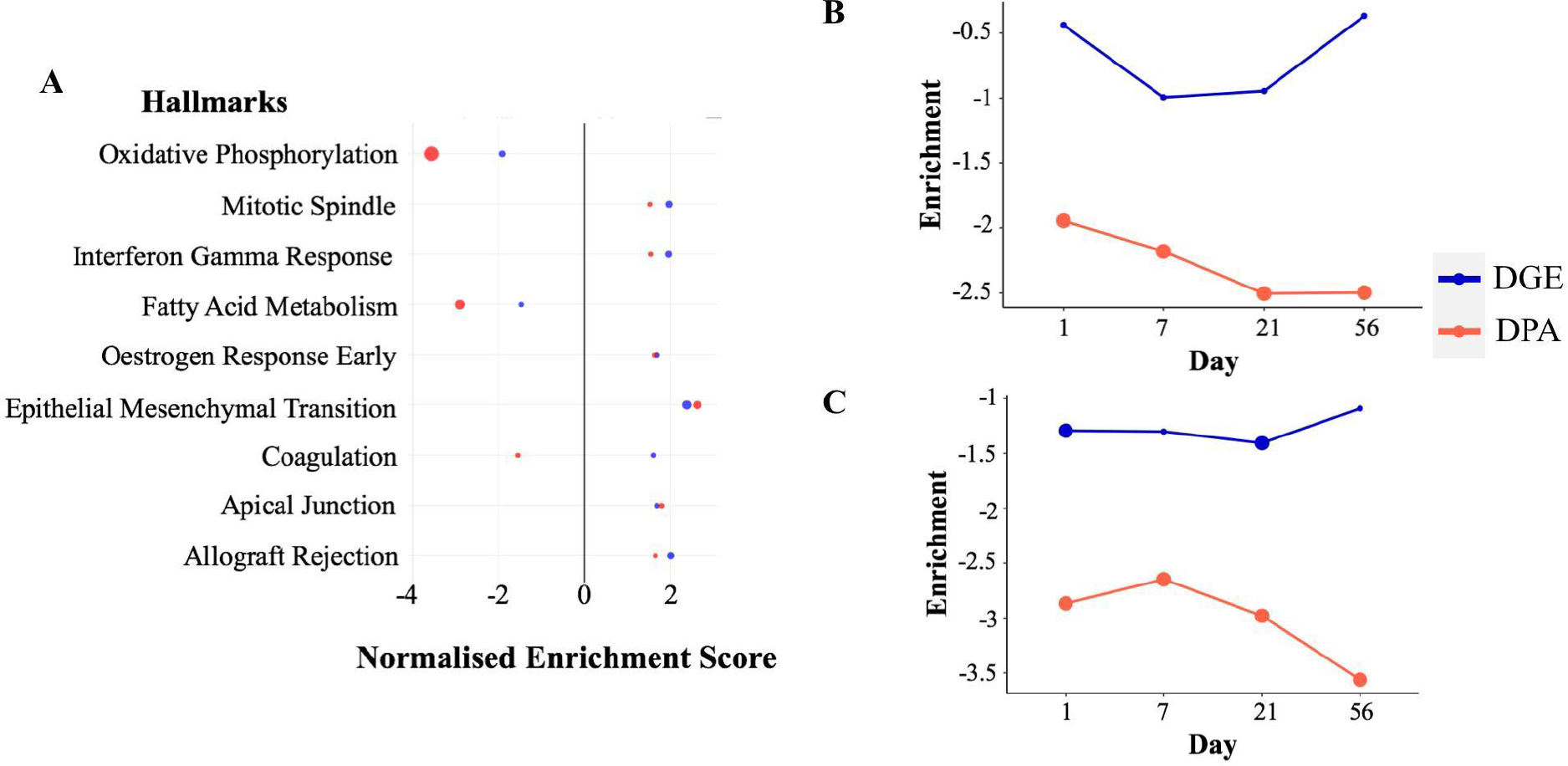
Enrichment scores from the differential gene (red dot) and protein (blue dot) expression analyses for the significantly regulated MSigDB Hallmarks sets in DGE and DPA (Panel A, TAC vs Sham). Time-resolved enrichment scores for ‘Fatty Acid Metabolism’ (Panel B) and ‘Oxidative Phosphorylation’ (Panel C) across time. Significant enrichment scores (pval<=0.1) have been highlighted with larger filled circles in the plot.

It can be observed that we have an altered regulation of fatty acid metabolism at the protein level starting at the early time-point Day1 which is reflected more predominantly at the latter time-point (Day21) (Figure 3B). Such regulation is of a negative sign when comparing the average expression of gene-set members for the TAC condition when compared to Sham, meaning that the above-mentioned gene-set is down-regulated. Similarly, we have a significant down-regulation of the ‘Oxidative Phosphorylation’ processes at both gene and protein levels, starting from the very early time-point (Day1). Another interesting observation is that the majority of gene members (56.94%) of the ‘Fatty Acid Metabolism’ and ‘Oxidative Phosphorylation’, seem to be associated with the *paleturquoise* cluster of genes that we have obtained from the clustering analysis with WGCNA. Gene members of such a cluster are shown to have a very strong and significant negative correlation with the TAC vs Sham comparisons (at all time points individually as well as combined) as well as a significant negative correlation with the HW/BW phenotype and a strong and significant positive correlation with the EF phenotype.

To conclude, these time-course analyses illustrate that changes in the expression of metabolic genes occur early in the development of pressure overload-induced heart failure before the detection of massive hypertrophy and contractile impairment. Of note, these findings are much in line with previous observations on early transcriptional alterations in the heart under chronic catecholamine exposure, indicating a general principle of metabolic gene regulation as an early response to chronic cardiac stress (31).

### Differential transcript usage

Another aspect of gene regulation involves alternative RNA splicing. Generally, intronic sequences get removed from pre-mRNA molecules during mRNA maturation. This process may also affect the combination of exons, which get included in the final mRNA product. In TACOMA, we have placed special attention on visualizing alternative splicing effects since they play a critical role in cardiovascular diseases by modulating gene expression and protein function, influencing processes such as heart muscle contraction and remodelling (32).

We performed enrichment analyses over all DTU results using Gene Ontology (BP ontology) and Hallmark gene sets from MSigDB (26) (Supplementary Figures 7-8) to identify biological processes and pathways that may be enriched for alternative splicing and could have been missed in a gene-level analysis. Our initial analysis revealed a distinct pattern of differential transcript usage between the TAC and Sham groups. For example, muscle contraction (GO:0006936), which consists of genes that are involved in generating force for muscle contraction, is one of the most significantly enriched gene sets for the main TAC vs Sham comparison (FDR = 0.0257). Another interesting significant term was Cell Cycle (GO:0007049) (FDR = 0.0072). Increased cell proliferation gene expression in adult cardiomyocytes has also been observed after TAC, the functional relevance in the adult heart however is not understood in detail (33). Further results pointed towards the regulation of gene isoforms involved in particular aspects of the cell cycle such as the G2 to M transition phase as witnessed by the statistically significant G2M Checkpoint hallmark set (FDR = 1.0223e-05), thus suggesting an enhanced proliferative activity in response to TAC (Figure 4).

**Figure 4:**
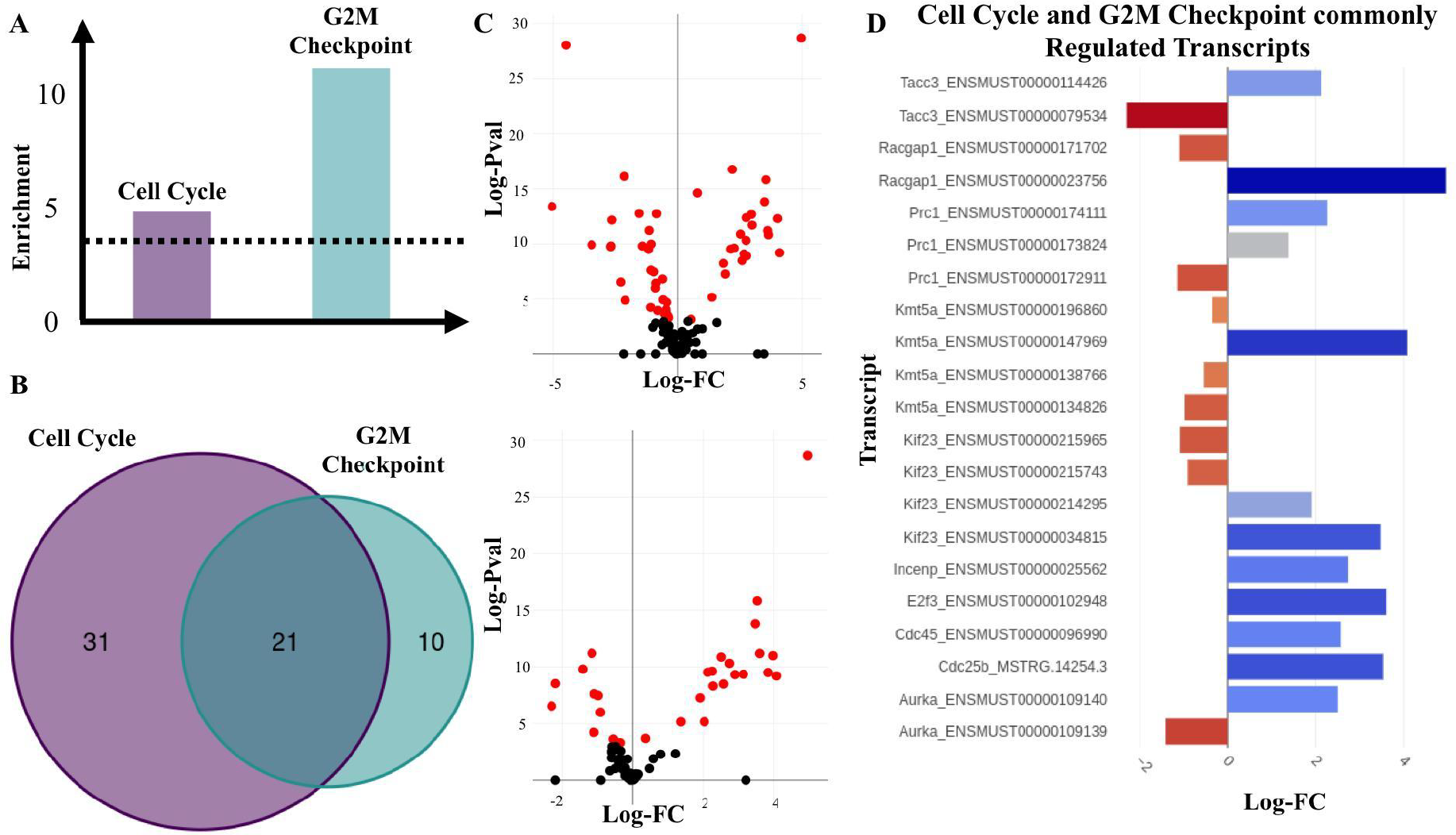
Transcripts of genes involved in ‘Cell Cycle’ and ‘G2M Checkpoint’ gene-sets. **A)** Enrichment scores of the ‘Cell Cycle’ and ‘G2M Checkpoint’ gene sets estimated as -Log (natural logarithm) enrichment of p-value significance scores (in dashed lines is shown the significance threshold - p.adj=0.05) **B)** Counts of significantly regulated transcripts of genes involved in the two gene-sets. **C)** Volcano plot of the DTU analysis for Cell Cycle (top) and G2M checkpoint (bottom). Red dots are significantly regulated transcripts, p.adj<=0.05). **D)** Significantly regulated transcripts (p.adj<=0.05) that are members of both the ‘Cell Cycle’ and ‘G2M Checkpoint’ gene sets.

Among the genes with significantly regulated transcript usage, we have identified Racgap1 (ENSMUST00000023756 and ENSMUST00000171702) and Kif23 (ENSMUST00000214295, ENSMUST00000215743 and ENSMUST00000215965) which are known to perform essential functions in central spindle formation (34). Interestingly, ‘Mitotic Spindle’ was also one of the Hallmark gene sets which appeared to have been significantly regulated in the DTU analyses for the TAC vs Sham comparison which is an event characteristic of cell division (35).

### Alternative usage of Tpm2 variants between RV and LV

Differential transcript usage can lead to the production of different protein isoforms from the same gene and may result in different functions of the RNA or protein product. Similar to our DTU analysis in the previous section, enrichment analyses over genes with significant changes in exon usage events were performed using Gene Ontology (BP ontology). So far, we have not reported on changes between the right (RV) and left (LV) ventricles of the heart following TAC. Our enrichment analysis for the RV vs LV comparison revealed two biological processes, which were associated with alternative RNA splicing: cardiac muscle contraction (GO:0060048) (FDR=0.00767). One of the most striking observations was the differential expression of transcript isoforms of the Tropomyosin 2-beta (Tpm2) gene, another known regulator of muscle contraction (Figure 5) which is shown to be commonly spliced in the heart (36).

**Figure 5:**
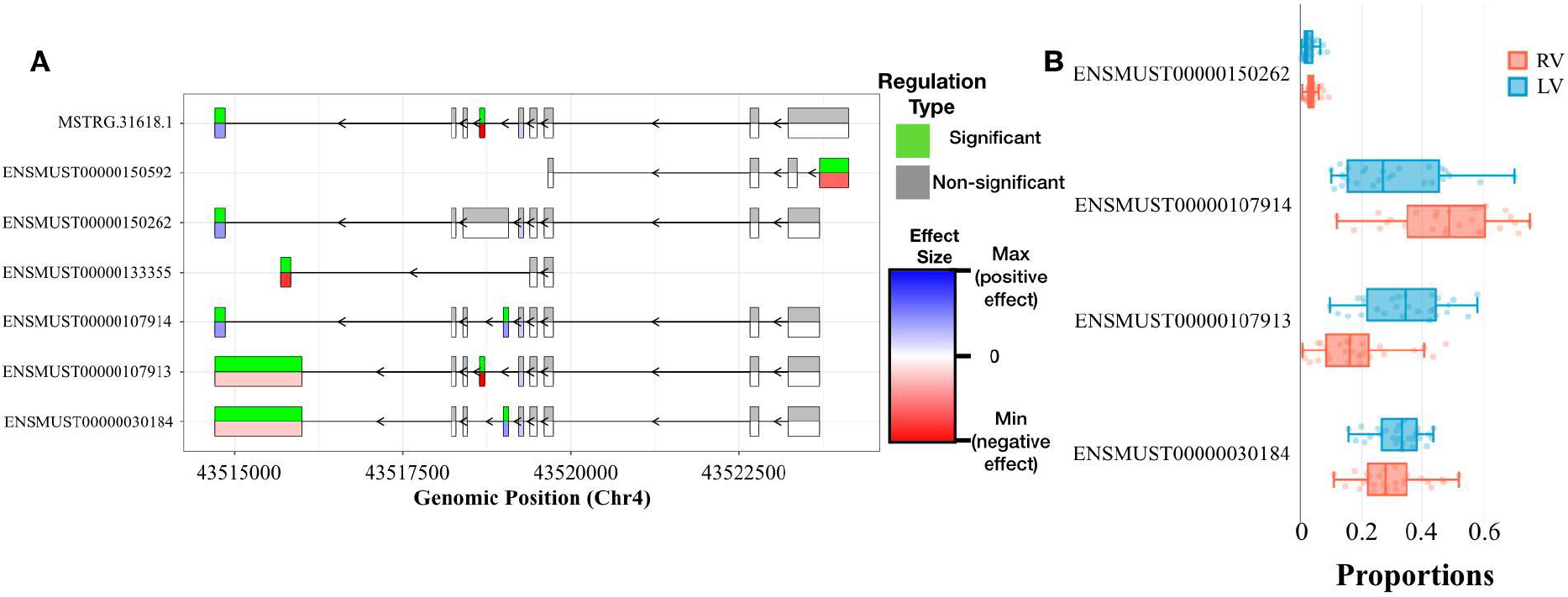
DEX and DTU Views of Tpm2 gene. **A)** Differential expression of individual exons of Tpm2 genes and transcripts for the RV vs LV comparisons. For each exon, the top half indicates whether the differential expression is significant (green) or not (grey), while the bottom half indicates the level of change (blue if we have a positive size effect and red for a negative size effect). **B)** Box plot showing the differences in usage between RV and LV of Tpm2 transcripts.

Figure 5A shows the Tpm2 exon usage pattern, which differs significantly between the RV and LV. Overall, we could identify six exons with an alternative usage pattern either pointing towards a preferential inclusion or exclusion in RV over LV. In Figure 5B it is provided additional details on the consequences with regard to transcript usage of robustly expressed transcripts. The changes in ENSMUST00000107913 and ENSMUST00000107914 are the only significant after correction for multiple testing. Both transcripts express two different proteins. While the first expresses a skeletal muscle isoform, the second produces a smooth-muscle isoform. Tropomyosin plays a crucial role in regulating the contraction process by facilitating the interaction between actin-containing thin filaments and myosin-containing thick filaments within muscles. In non-muscle cells expressing various tropomyosin isoforms, tropomyosins are actively involved in numerous cellular events related to the cytoskeleton. These findings suggest a selective upregulation of one specific Tpm2 isoform in the RV (ENSMUST00000107914), potentially contributing to the differential contractile response of the ventricles under TAC-induced stress. To the best of our knowledge, the functional implications of the two protein isoforms of Tpm2 are not fully understood yet.

### Novel Genes

Upon de-novo assembly of the GRCm38.102 reference genome with long-reads Nanopore cDNA transcriptomics with StringTie2 (v2.2.1), we identified 84 genes with completely novel transcripts i.e. no overlap with any annotated gene. Of these 84 genes, 33 of them were significant in at least one of the DGE comparisons that we have tested. Upon the identification of the novel genes, we then performed open reading frame (ORF) identification over each novel gene sequence by using the *findORFs()* function from the ORFik R-package (v1.20.2, citation needed). The ORF DNA sequences were then translated into protein sequences by using the *translate()* function from Biostrings R-package (v2.68.1) which was followed up by domain annotation analysis with Interproscan (v5.65-97.0) (14). From such an analysis, Interproscan was able to predict functional domains for 26 out of 84 novel genes which were significant in at least one of the DGE comparisons.

## Discussions

TACOMA enables interactive online analysis, exploration, integration, and visualisation of a new multi-omics time course dataset from a TAC mouse model. To the best of our knowledge, there are no interactive web-applications for integrated proteomics and transcriptomics data exploration in the cardiovascular field. However, the integration of various gene expression datasets in heart failure was recently addressed by the ReHeat (37) and the Magnetique (27) portals, which are also available as online applications. ReHeat comprises a comprehensive meta-analysis of public human HF microarray and RNA-seq datasets, while Magnetique used mRNA-seq data from the Myocardial Applied Genomics Network (MAGNet) consortium. While ReHeat focused on the analysis at the gene level, Magnetique added special attention to the analysis at the transcript level by providing differential RNA transcript isoform usage (DTU) changes and predicting RNA-binding protein (RBP) to target transcript interactions using a Global test approach.

TACOMA goes beyond a simple exploration of a new multi-omics data-set by identifying clusters of co-expressed genes and putting special emphasis on exon- and transcript-level analysis. Evidently, the interplay between the proteomics and transcriptomics layers is well represented as well. Additionally, we enhanced the known cardiac transcriptome by a *de novo* assembly, which we obtained from Illumina and cDNA Nanopore reads.

Several known and novel findings have been presented by example. First, we reported on a notable shift in energy metabolism within hypertrophied hearts, transitioning from fatty acid metabolism to glucose and glycolysis. This metabolic shift was initiated at the gene level by Day 7 and completed by Day 21 after TAC, as evidenced by the significant regulatory patterns in gene and protein expressions associated with fatty acid oxidation. Similar to fatty acids, through TACOMA, we were able to demonstrate a significant down-regulation of the tricarboxylic acid (TCA) cycle gene set in cardiac tissue post-TAC, particularly evident at week 8, suggesting a link between TCA cycle disruption and the progression to heart failure, corroborated by consistent gene and protein expression patterns. Second, through TACOMA we were able to identify differential exon-skipping events in key cardiac genes, notably in Tpm2 isoforms, between the right and left ventricles post-TAC, suggesting contractile differences and providing potential new insights into the molecular mechanisms of heart contraction under stress.

In the future, we will expand TACOMA in terms of new functionalities and data sets. Additional animal models of heart disease will be added and several other functional analysis methods of multi-omics data analysis will be included.

## Conclusions

In this study, we have produced and analysed a comprehensive time-series proteomics and transcriptomics data set from a TAC mouse model and provided subsequent analysis results through the TACOMA web-application (https://shiny.dieterichlab.org/app/tacoma). The design included three factors (TAC vs Sham, time and LV vs RV) and we paid special attention to the details of the statistical modelling. TACOMA is unique in integrating proteomics and transcriptomics data for a pressure-overload mouse model of heart failure in a user-friendly web application. We anticipate that TACOMA will be adopted by clinician scientists and cardiovascular research as an exploratory tool to further uncover relevant molecular mechanisms associated with HF progression and/or to make comparisons with their own independent studies. Future work about TACOMA will focus on the addition of other functional analysis methods and more layers of omes. We also plan to open up TACOMA to integrate private data from users through authentication-based mechanisms.

## Acknowledgements

We thank Harald Wilhelmi for excellent support in setting up the computational framework as well as for excellent technical assistance and Etienne Boileau for fruitful discussions.

## References

1. Wang X., Zhu X., Shi L., Wang J., Xu Q., Yu B., et al. (2023) A time-series minimally invasive transverse aortic constriction mouse model for pressure overload-induced cardiac remodeling and heart failure. Front Cardiovasc Med. 2023;10: 1110032.

2. Xia Y., Lee K., Li N., Corbett D., Mendoza L., Frangogiannis N.G. (2008) Characterization of the inflammatory and fibrotic response in a mouse model of cardiac pressure overload. Histochem Cell Biol. 2008;131: 471–481.

3. Roehr J.T., Dieterich C., Reinert K. (2017) Flexbar 3.0 – SIMD and multicore parallelization. Bioinformatics. 2017;33: 2941–2942.

4. Langmead B., Salzberg S.L. (2012) Fast gapped-read alignment with Bowtie 2. Nat Methods. 2012;9: 357–359.

5. Dobin A., Davis C.A., Schlesinger F., Drenkow J., Zaleski C., Jha S., et al (2012). STAR: ultrafast universal RNA-seq aligner. Bioinformatics. 2012;29: 15–21.

6. Kovaka S., Zimin A.V., Pertea G.M., Razaghi R., Salzberg S.L., Pertea M. (2019) Transcriptome assembly from long-read RNA-seq alignments with StringTie2. Genome Biol. 2019;20: 278.

7. Li H. (2018) Minimap2: pairwise alignment for nucleotide sequences. Bioinformatics. 2018;34: 3094–3100.

8. Pertea G., Pertea M. (2020) GFF Utilities: GffRead and GffCompare. F1000Res. 2020;9: 304.

9. Patro R., Duggal G., Love M.I., Irizarry R.A., Kingsford C. (2017) Salmon provides fast and bias-aware quantification of transcript expression. Nature Methods. volume 14, pages 417–419.

10. Love M.I., Huber W., Anders S. (2014) Moderated estimation of fold change and dispersion for RNA-seq data with DESeq2. Genome Biology volume 15, Article number: 550.

11. Robinson M.D., McCarthy D.J., Smyth G.K. (2010) edgeR: a Bioconductor package for differential expression analysis of digital gene expression data. Bioinformatics. 2010;26: 139–140.

12. Durinck S., Spellman P.T., Birney E., Huber W. (2009) Mapping identifiers for the integration of genomic datasets with the R/Bioconductor package biomaRt. Nat Protoc. 2009;4: 1184–1191.

13. Tjeldnes H., Labun K., Torres C.Y., Chyżyńska K., Świrski M., Valen E. (2021) ORFik: a comprehensive R toolkit for the analysis of translation. BMC Bioinformatics. 2021;22: 336.

14. Blum M., Chang H.Y., Chuguransky S., Grego T., Kandasaamy S., Mitchell A., et al. (2021) The InterPro protein families and domains database: 20 years on. Nucleic Acids Res. 2021;49: D344–D354.

15. Boileau E., Doroudgar S., Riechert E., Jürgensen L., Ho T.C., Katus H.A., et al. (2020) A Multi-Network Comparative Analysis of Transcriptome and Translatome Identifies Novel Hub Genes in Cardiac Remodeling. Front Genet. 2020;11: 583124.

16. Langfelder P., Horvath S. (2012) Fast R functions for robust correlations and hierarchical clustering. J Stat Softw. 2012;46. doi:10.18637/jss.v046.i11

17. Hughes C.S., Foehr S., Garfield D.A., Furlong E.E., Steinmetz L.M., Krijgsveld J. (2014) Ultrasensitive proteome analysis using paramagnetic bead technology. Mol Syst Biol. 2014;10: 757.

18. Hughes C.S., Moggridge S., Müller T., Sorensen P.H., Morin G.B., Krijgsveld J. (2019) Single-pot, solid-phase-enhanced sample preparation for proteomics experiments. Nat Protoc. 2019;14: 68–85.

19. Werner T., Sweetman G., Savitski M.F., Mathieson T., Bantscheff M., Savitski M.M. (2014) Ion coalescence of neutron encoded TMT 10-plex reporter ions. Anal Chem. 2014;86: 3594–3601.

20. Reichel M., Liao Y., Rettel M., Ragan C., Evers M., Alleaume A-M., et al. (2016) In Planta Determination of the mRNA-Binding Proteome of Arabidopsis Etiolated Seedlings. Plant Cell. 2016;28: 2435–2452.

21. Franken H., Mathieson T., Childs D., Sweetman G.M.A., Werner T., Tögel I., et al. (2015) Thermal proteome profiling for unbiased identification of direct and indirect drug targets using multiplexed quantitative mass spectrometry. Nat Protoc. 2015;10: 1567–1593.

22. Feng Z., Fang P., Zheng H., Zhang X. (2023) DEP2: an upgraded comprehensive analysis toolkit for quantitative proteomics data. Bioinformatics. 2023;39: btad526.

23. Ritchie M.E., Phipson B., Wu D., Hu Y., Law C.W., Shi W., et al. (2015) limma powers differential expression analyses for RNA-sequencing and microarray studies. Nucleic Acids Res. 2015;43: e47.

24. Strimmer K. (2008) A unified approach to false discovery rate estimation. BMC Bioinformatics. 2008;9: 303.

25. Korotkevich G., Sukhov V., Budin N., Shpak B., Artyomov M.N., Sergushichev A. (2021) Fast gene set enrichment analysis. bioRxiv. 2021. p. 060012. doi:10.1101/060012

26. Subramanian A., Tamayo P., Mootha V.K., Mukherjee S., Ebert B.L., Gillette M.A., et al. (2005) Gene set enrichment analysis: a knowledge-based approach for interpreting genome-wide expression profiles. Proc Natl Acad Sci U S A. 2005;102: 15545–15550.

27. Britto-Borges T., Ludt A., Boileau E., Gjerga E., Marini F., Dieterich C. (2022) Magnetique: an interactive web application to explore transcriptome signatures of heart failure. J Transl Med. 2022;20: 513.

28. Gustavsson E.K., Zhang D., Reynolds R.H., Garcia-Ruiz S., Ryten M. (2022) ggtranscript: an R package for the visualization and interpretation of transcript isoforms using ggplot2. Bioinformatics. 2022;38: 3844–3846.

29. Lopaschuk G.D., Karwi Q.G., Tian R., Wende A.R., Dale Abel E. (2021) Cardiac Energy Metabolism in Heart Failure. Circ Res. 2021. doi:10.1161/CIRCRESAHA.121.318241

30. Bertero E., Maack C. (2018) Metabolic remodelling in heart failure. Nat Rev Cardiol. 2018;15: 457–470.

31. Dewenter M., Pan J., Knödler L., Tzschöckel N., Henrich J., Cordero J., et al. (2022) Chronic isoprenaline/phenylephrine vs. exclusive isoprenaline stimulation in mice: critical contribution of alpha1-adrenoceptors to early cardiac stress responses. Basic Res Cardiol. 2022;117: 1–23.

32. Regulation of splicing in cardiovascular disease. Epigenetics in Cardiovascular Disease. Academic Press; 2021. pp. 163–186.

33. Froese N., Cordero J., Abouissa A., Trogisch F.A., Grein S., Szaroszyk M., et al. (2022) Analysis of myocardial cellular gene expression during pressure overload reveals matrix based functional intercellular communication. iScience. 2022;25: 103965.

34. Coudert E., Gehant S., de Castro E., Pozzato M., Baratin D., Neto T., et al. (2023) Annotation of biologically relevant ligands in UniProtKB using ChEBI. Bioinformatics. 2023;39. doi:10.1093/bioinformatics/btac793

35. Drazen J.M. (2001) Expression of Concern: Beltrami A.P. et al. Evidence That Human Cardiac Myocytes Divide after Myocardial Infarction. N Engl J Med 2001;344:1750–7 and Quaini F et al. Chimerism of the Transplanted Heart. N Engl J Med 2002;346:5–15. N Engl J Med. 2018;379: 1870.

36. Montañés-Agudo P., Pinto Y.M., Creemers E.E. (2023) Splicing factors in the heart: Uncovering shared and unique targets. J Mol Cell Cardiol. 2023;179: 72–79.

37. Ramirez Flores R.O., Lanzer J.D., Holland C.H., Leuschner F., Most P., Schultz J., et al. (2021) Consensus Transcriptional Landscape of Human End-Stage Heart Failure. J Am Heart Assoc. 2021. doi:10.1161/JAHA.120.019667

