## Supplementary Notes 1 for "Transverse Aortic COnstriction Multi-omics Analysis (TACOMA) uncovers pathophysiological cardiac molecular mechanisms"

Tables of comparisons performed for the DGE, DXE and DTU analysis and their contrast coefficients from the defined design matrix model.matrix(~ Ventricle + Day + Condition + Ventricle:Condition + Day:Condition, data=metatada) are provided below:

| | Condition | Position | Day | Pos:Cond | Day:Cond | 7 vs 1 | 21 vs 7 | 56 vs 21 | TAC_1 | TAC_7 | TAC_21 | TAC_56 |
|---|---|---|---|---|---|---|---|---|---|---|---|---|
| **Intercept** | 0 | 0 | 0 | 0 | 0 | 0 | 0 | 0 | 0 | 0 | 0 | 0 |
| **PositionRV** | 0 | 1 | 0 | -1 | 0 | 0 | 0 | 0 | 0 | 0 | 0 | 0 |
| **Day1** | 0 | 0 | 0.5 | 0 | 0 | -1 | 0 | 0 | -1 | 0 | 0 | 0 |
| **Day7** | 0 | 0 | 0.5 | 0 | 0 | 1 | -1 | 0 | 0 | -1 | 0 | 0 |
| **Day21** | 0 | 0 | 0.5 | 0 | 0 | 0 | 1 | -1 | 0 | 0 | -1 | 0 |
| **Day56** | 0 | 0 | 0.5 | 0 | 0 | 0 | 0 | 1 | 0 | 0 | 0 | -1 |
| **ConditionTAC** | 1 | 0 | 0 | 0 | 0 | 0 | 0 | 0 | 0 | 0 | 0 | 0 |
| **PosRV:CondTAC** | 0.5 | 0.5 | 0 | 1 | 0 | 0 | 0 | 0 | 0 | 0 | 0 | 0 |
| **Day1:CondTAC** | 0.25 | 0 | 0.25 | 0 | 0.25 | -0.5 | 0 | 0 | 1 | 0 | 0 | 0 |
| **Day7:CondTAC** | 0.25 | 0 | 0.25 | 0 | 0.25 | 0.5 | -0.5 | 0 | 0 | 1 | 0 | 0 |
| **Day21:CondTAC** | 0.25 | 0 | 0.25 | 0 | 0.25 | 0 | 0.5 | -0.5 | 0 | 0 | 1 | 0 |
| **Day56:CondTAC** | 0.25 | 0 | 0.25 | 0 | 0.25 | 0 | 0 | 0.5 | 0 | 0 | 0 | 1 |

The questions being addressed as well as the comparisons being performed can be described as below:

| Question | Comparison |
| --- | --- |
| *Which genes change by condition?* | Condition: The main differential gene expression comparison is evaluating the difference between the TAC and Sham conditions, but with additional weighting on the interactions of 'Condition' with the 'Day' and 'Position' terms. |
| *Which genes change by ventricle?* | Ventricle: The main differential gene expression comparison is evaluating the difference between RV and LV positions, but with additional weighting on the interaction between the 'Position' and the 'Condition' terms. |
| *Which genes change at all time-points compared to control?* | Day: This differential gene expression comparison evaluates the combination of the main effect 'Day' and their interaction with the 'Condition' term. |
| *Which genes change in any day?* | Day.Condition: Here, we seek to test the effect of interaction between the 'Day' and the 'Condition' terms by weighting them equaly. |
| *Which genes change in TAC for a day?* | Day1.ConditionvsDay1: Here, we test the difference in gene expression between samples taken on Day1 and those taken on Day1 after being subjected to the ConditionTAC. The comparison aims to identify the differential gene expression due to the TAC condition specifically on Day 1. |
| | Day7.ConditionvsDay7: Here, we test the difference in gene expression between samples taken on Day7 and those taken on Day7 after being subjected to the ConditionTAC. The comparison aims to identify the differential gene expression due to the TAC condition specifically on Day 7. |
| | Day21.ConditionvsDay21: Here, we test the difference in gene expression between samples taken on Day21 and those taken on Day21 after being subjected to the ConditionTAC. The comparison aims to identify the differential gene expression due to the TAC condition specifically on Day 21. |

| | |
|---|---|
| | Day56.ConditionvsDay56: Here, we test the difference in gene expression between samples taken on Day56 and those taken on Day56 after being subjected to the ConditionTAC. The comparison aims to identify the differential gene expression due to the TAC condition specifically on Day 56. |
| *Which genes change in TAC for ventricle?* | Ventricle.Condition: Here, we are testing for genes that have a differential expression due to the combination of being in the RV position and being subjected to the TAC condition, relative to just being in the RV position alone. |
| *Which genes change over time?* | Day7vsDay1: Here, we are testing for genes that for genes whose expression changes between Day7 and Day1 and to assess how this change is influenced when the ConditionTAC. |
| | Day21vsDay7: Here, we are testing for genes that for genes whose expression changes between Day21 and Day7 and to assess how this change is influenced when the ConditionTAC. |
| | Day56vsDay21: Here, we are testing for genes that for genes whose expression changes between Day56 and Day21 and to assess how this change is influenced when the ConditionTAC. |

### Supplementary Figures

#### Supplementary Figure 1

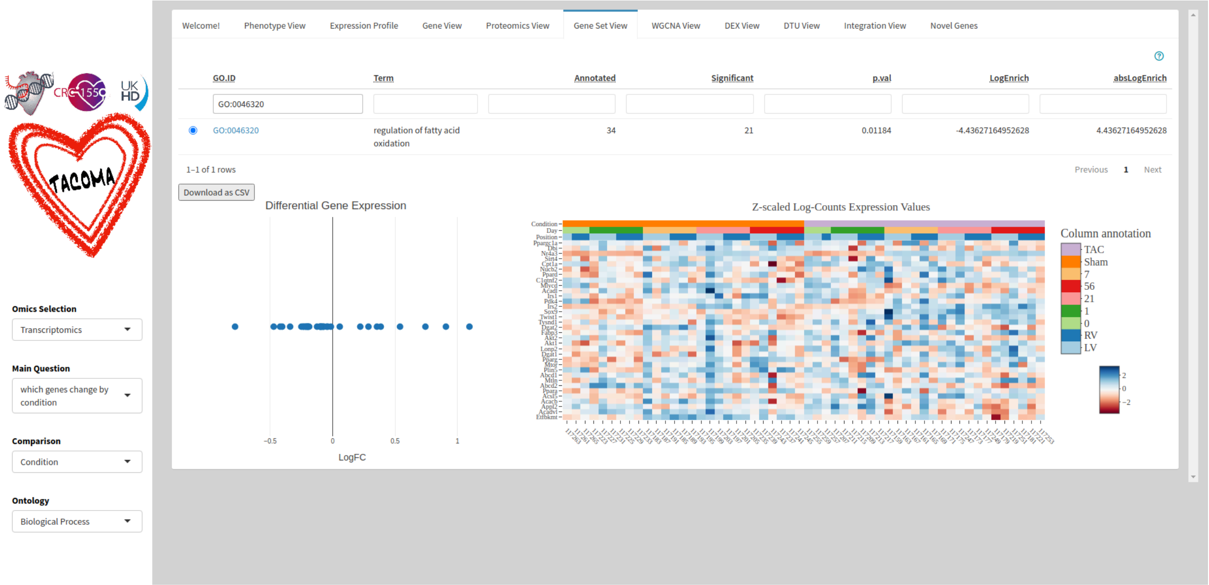


Gene expression values and enrichment scores (LogEnrich) based on the differential gene expression analysis for the *"regulation of fatty acid oxidation"* (GO:0046320) gene-set members as visualized in TACOMA.

#### Supplementary Figure 2

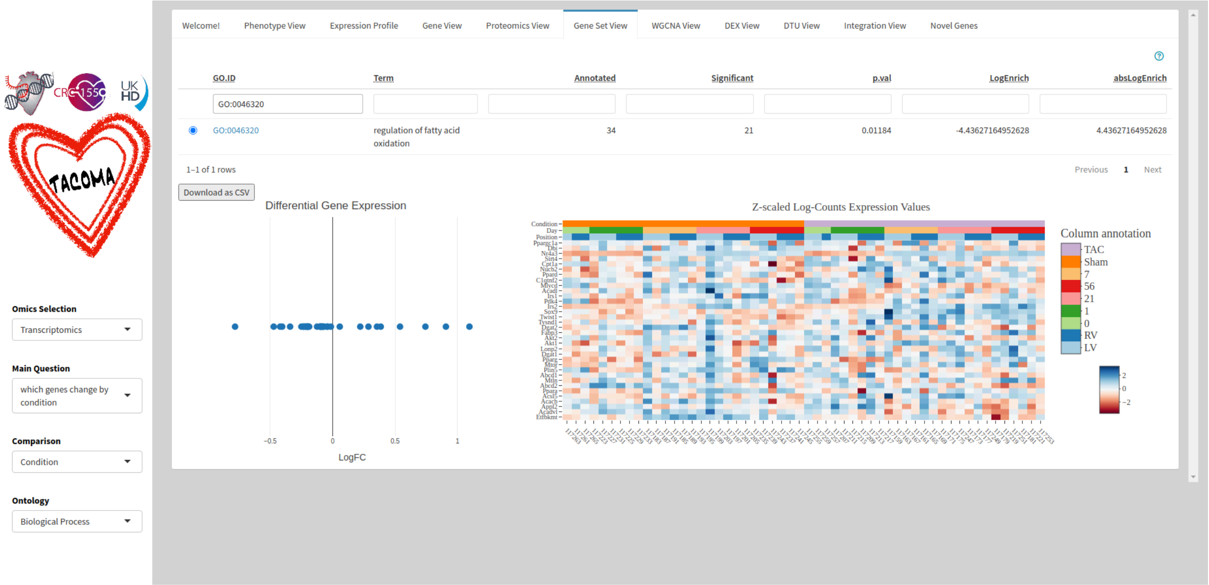


Protein abundance values and enrichment scores (LogEnrich) based on the differential protein abundance analysis for the *"regulation of fatty acid oxidation"* (GO:0046320) gene-set members as visualized in TACOMA.

### Supplementary Figure 3

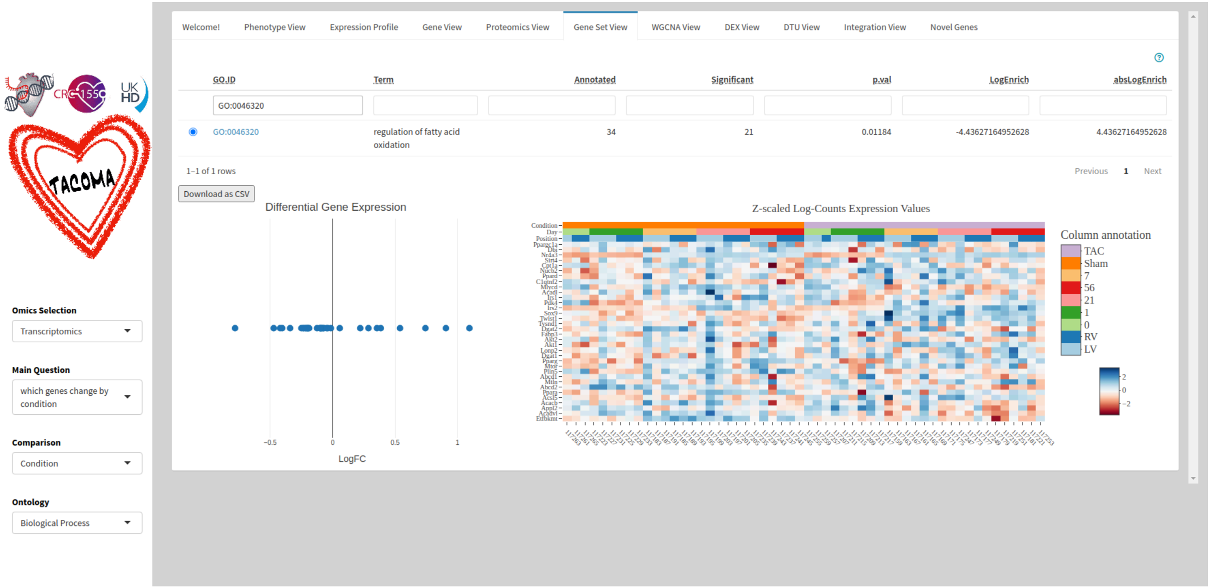


Gene expression values and enrichment scores (LogEnrich) based on the differential gene expression analysis for the *"oxidative phosphorylation"* (GO:0006119) gene-set members as visualized in TACOMA.

### Supplementary Figure 4

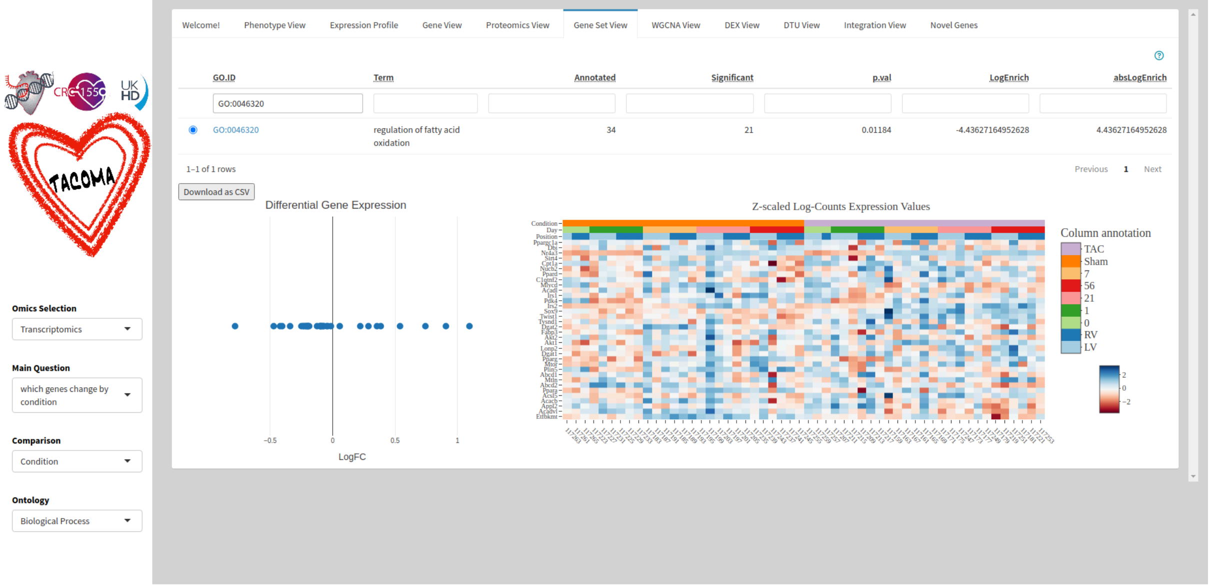


Protein abundance values and enrichment scores (LogEnrich) based on the differential protein abundance analysis for the *"oxidative phosphorylation"* (GO:0006119) gene-set members as visualized in TACOMA.

### Supplementary Figure 5

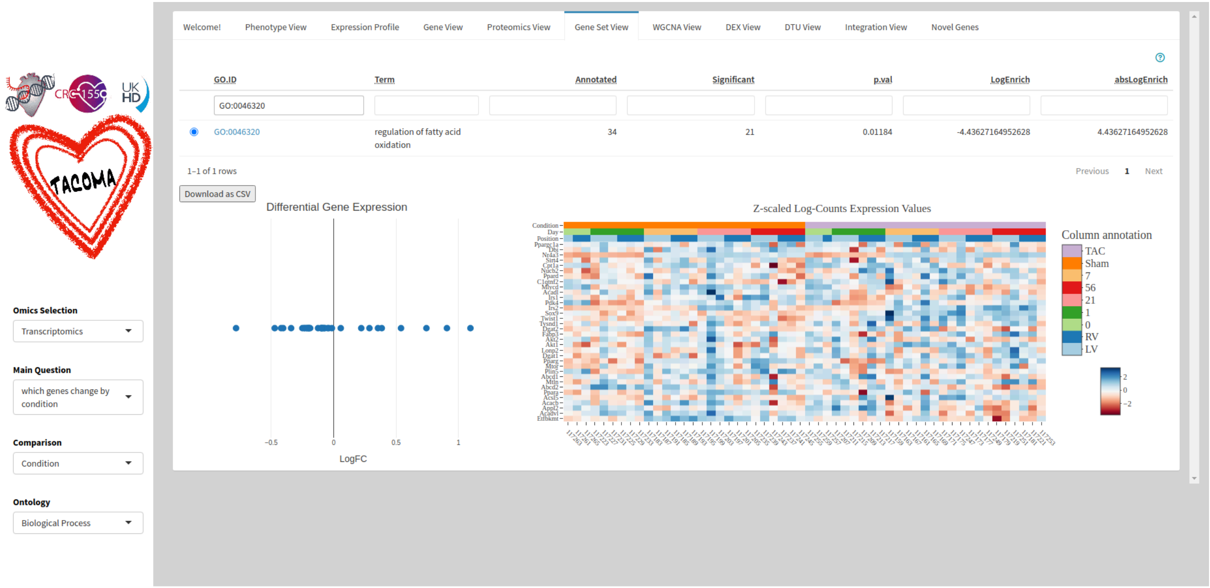


Protein abundance values and enrichment scores (LogEnrich) based on the differential protein abundance analysis for the *"tricarboxylic acid cycle"* (GO:0006099) gene-set members as visualized in TACOMA.

### Supplementary Figure 6

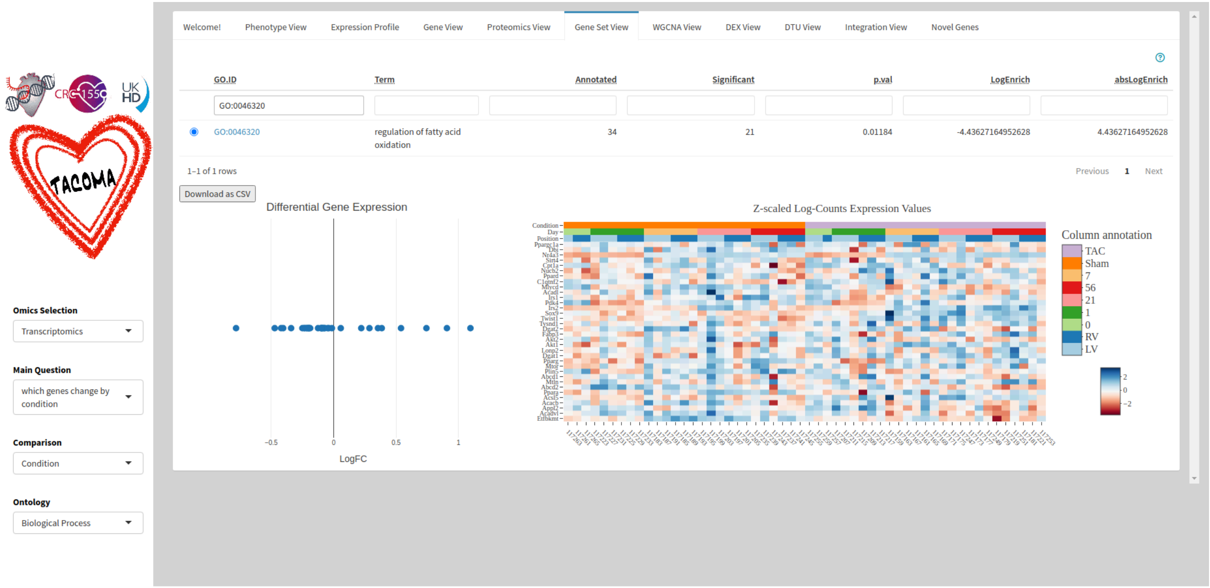


Gene expression values and enrichment scores (LogEnrich) based on the differential gene expression analysis for the *"tricarboxylic acid cycle"* (GO:0006099) gene-set members as visualized in TACOMA.

### Supplementary Figure 7

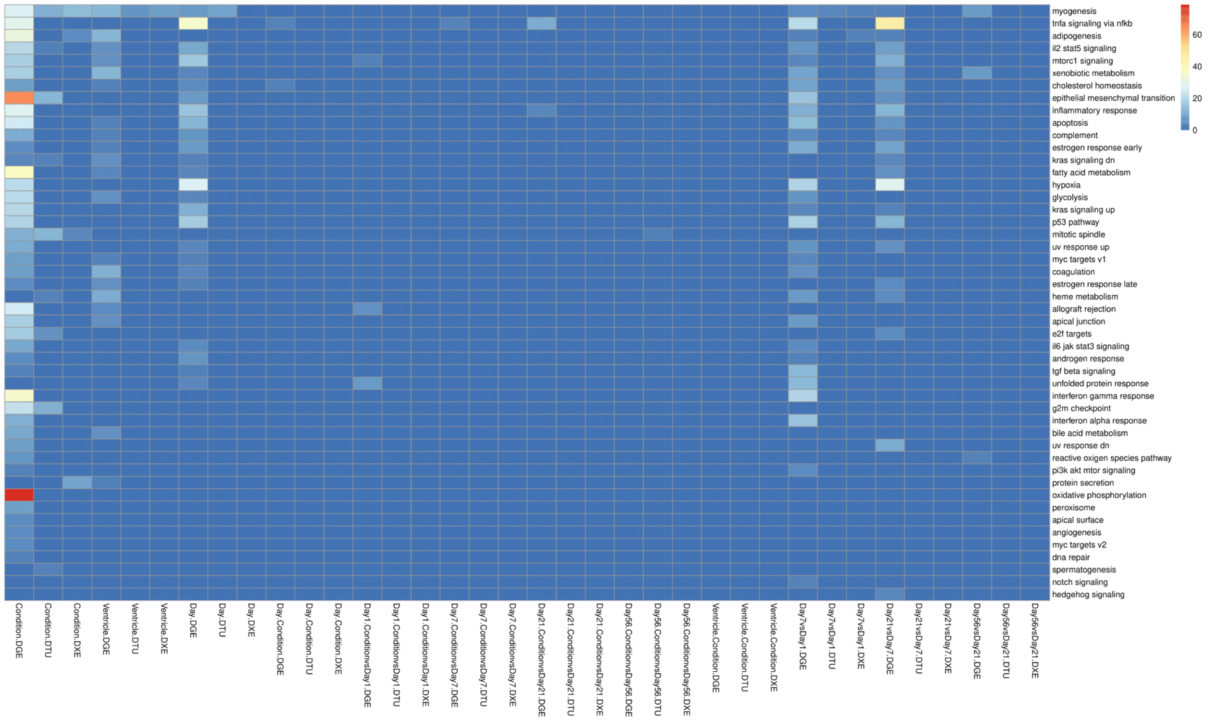


Enrichment scores (LogEnrich) of Gene Ontology (BP ontology) sets based on the DGE, DTU and DXE differential results for all the 12 comparisons performed. Boxes with colors as displayed on the color legend to the left of the heatmap, correspond to significant enrichment cases (p.adj <= 0.05).

### Supplementary Figure 8

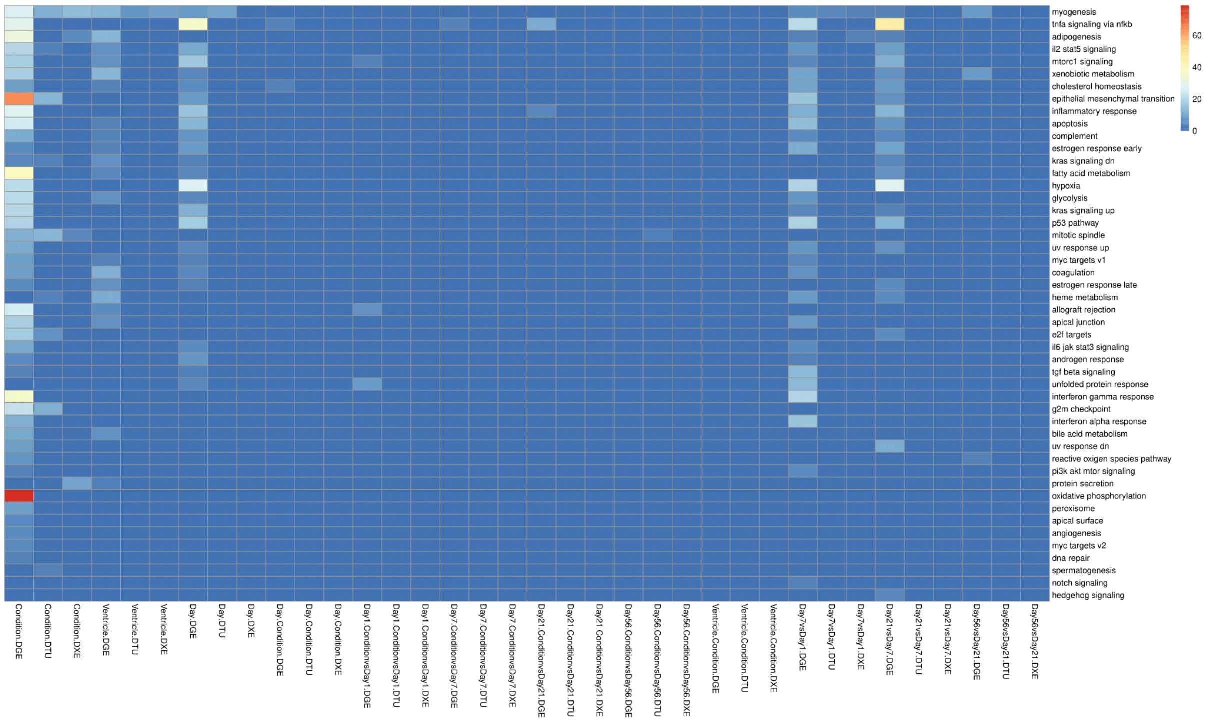


Enrichment scores (LogEnrich) of Hallmark gene sets from MSigDB based on the DGE, DTU and DXE differential results for all the 12 comparisons performed. Boxes with colors as displayed on the color legend to the left of the heatmap, correspond to significant enrichment cases (p.adj <= 0.05).
